# gen3sis: the <u>gen</u>eral <u>e</u>ngine for <u>e</u>co-<u>e</u>volutionary <u>si</u>mulation<u>s</u> on the origins of biodiversity

**DOI:** 10.1101/2021.03.24.436109

**Authors:** Oskar Hagen, Benjamin Flück, Fabian Fopp, Juliano S. Cabral, Florian Hartig, Mikael Pontarp, Thiago F. Rangel, Loïc Pellissier

## Abstract

Understanding the origins of biodiversity has been an aspiration since the days of early naturalists. The immense complexity of ecological, evolutionary and spatial processes, however, has made this goal elusive to this day. Computer models serve progress in many scientific fields, but in the fields of macroecology and macroevolution, eco-evolutionary models are comparatively less developed. We present a general, spatially-explicit, eco-evolutionary engine with a modular implementation that enables the modelling of multiple macroecological and macroevolutionary processes and feedbacks across representative spatio-temporally dynamic landscapes. Modelled processes can include environmental filtering, biotic interactions, dispersal, speciation and evolution of ecological traits. Commonly observed biodiversity patterns, such as α, β and γ diversity, species ranges, ecological traits and phylogenies, emerge as simulations proceed. As a case study, we examined alternative hypotheses expected to have shaped the latitudinal diversity gradient (LDG) during the Earth’s Cenozoic era. We found that a carrying capacity linked with energy was the only model variant that could simultaneously produce a realistic LDG, species range size frequencies, and phylogenetic tree balance. The model engine is open source and available as an R-package, enabling future exploration of various landscapes and biological processes, while outputs can be linked with a variety of empirical biodiversity patterns. This work represents a step towards a numeric and mechanistic understanding of the physical and biological processes that shape Earth’s biodiversity.

## Introduction

Ecological and evolutionary processes have created various patterns of diversity in living organisms across the globe [1]. Species richness varies across regions, such as continents [2, 3], and along spatial and environmental gradients [4, 5], such as latitude [6, 7]. These well-known patterns, derived from the observed multitude of life forms on Earth, have intrigued naturalists for centuries [1, 8, 9] and stimulated the formulation of numerous hypotheses to explain their origin [e.g. 1, 6, 7, 10, 11-15]. Ecologists and evolutionary biologists have attempted to test and disentangle these hypotheses [16], for example via models of cladogenesis [17] or correlative spatial analyses [18, 19]. However, to this day, a mechanistic understanding of ecological, evolutionary and geodynamical spatial dynamics driving diversity patterns remains elusive [20, 21].

The complexity of interacting ecological, evolutionary and spatial processes limits our ability to formulate, test and apply the mechanisms forming biodiversity patterns [22, 23]. Additionally, multiple processes act and interact with different relative strengths across spatio-temporal scales [20]. Current research suggests that allopatric [24–26] and ecological [22] speciation, dispersal [27] and adaptation [28] all act conjointly in interaction with the environment [29, 30], producing observed biodiversity patterns [31]. Comprehensive explanations of the origin and dynamics of biodiversity must therefore consider a large number of biological processes and feedbacks [32], including species’ ecological and evolutionary responses to their dynamic abiotic environment, acting on both ecological and evolutionary time scales [20, 33]. Consequently, biodiversity patterns can rarely be explained by a single hypothesis, as the expectations of the various contending mechanisms are not clearly asserted [20, 34].

A decade ago, a seminal paper by Gotelli and colleagues [35] formulated the goal of developing a “general simulation model for macroecology and macroevolution” (hereafter computer models). Since then, many authors have reiterated this call for a broader use of computer models in biodiversity research [20, 36, 37]. With computer models, researchers can explore with simulations the implications of implemented hypotheses and mechanisms and evaluate whether emerging simulated patterns are compatible with observations. Several case studies have illustrated the feasibility and usefulness of computer models in guiding intuition for the interpretation of empirical data [24, 26, 38–42]. Moreover, models have reproduced realistic large-scale biodiversity patterns, such as along latitude [25, 43, 44], by considering climate and geological dynamics [24, 26, 42], and population isolation by considering dispersal ability and geographic distance [24-26, 38-42]. For example, computer models were used to examine how oceans’ paleogeography influenced biodiversity dynamics in marine ecosystems [24, 41–43]. Nevertheless, the potential of computer models to enlighten the mechanisms underlying biodiversity patterns remains largely untapped.

Macroevolutionary studies have highlighted that patterns emerging from simulations are generally sensitive to the mechanisms implemented, and to the landscapes upon which those act [24, 25, 42, 43]. Systematically comparing and exploring the effects of mechanisms and landscapes, however, is often hindered by the lack of flexibility and idiosyncrasies of existing models. Most models implement, and thus test, only a limited set of evolutionary processes and hypotheses. Many models are designed for specific and therefore fixed purposes including spatial and temporal boundaries, ranging from the global [24, 25] to continental [26] or regional scale [39, 40], and in time, from millions of years [39, 40, 42, 43] to thousands of years [25, 26]. Moreover, previous eco-evolutionary population models were developed to test a fixed number of mechanisms [24, 25, 35, 40, 42, 44–50]. The diverse input and output formats and limited code availability [51], as well as the different algorithmic implementations, have reduced interoperability between hitherto available models. Biological hypotheses and landscapes should be compared within a common and standardized platform with the modularity required for flexible explorations of multiple landscapes and processes [35]. Increased generality is thus a desirable feature of computer models that aim to explore the mechanisms and landscapes that shape biodiversity in dynamic systems such as rivers [52], oceans [41, 42], islands [39, 40, 53] and mountains [54, 55], or across gradients such as latitude [20, 25, 43].

Here, we present a modelling engine that offers the possibility to explore eco-evolutionary dynamics of lineages under a broad range of biological processes and landscapes. Simulated species populations occupy a spatial domain (hereafter site) bounded by a combination of geological, climatic and ecological factors. The sites occupied by a species define the species’ realized geographic range (hereafter species range) [56]. The engine then tracks species populations over time, which can change as a result of dynamic environments, as well as species dispersal ability, ecological interactions, local adaptation and speciation. The initial species range and the criteria for speciation, dispersal, ecological interactions and trait evolution are adjustable mechanisms, allowing the integration of a wide range of hypotheses within the model. Given the flexibility of modifying both mechanisms and landscapes, we consider the engine a general tool and named it “<u>gen</u>eral <u>e</u>ngine for <u>e</u>co-<u>e</u>volutionary <u>si</u>mulation<u>s</u>” (hereafter gen3sis). We highlight the potential of gen3sis as a flexible tool to gain inferences about the underlying processes behind biodiversity patterns by tackling a long-standing topic in evolutionary ecology: the latitudinal diversity gradient (LDG) [20]. We implement three alternative hypotheses proposed to explain the LDG [20]: (i) *time for species accumulation* [57–60], (ii) *diversification rates* i.e. depending on temperature [61, 62], and *ecological limits* i.e. depending on energetic carrying capacity [63, 64]. We compare simulation results to empirical distribution and phylogenetic patterns of major tetrapod clades (i.e. mammals, birds, amphibians and reptiles).

## Engine principles and scope

Gen3sis is a modelling engine, developed for formalizing and testing multiple hypotheses about the emergence of biodiversity patterns. The engine simulates the consequences of multiple customizable processes and landscapes responsible for the appearance (speciation) and disappearance (extinction) of species over evolutionary time scales. Speciation and extinction emerge from ecological and evolutionary mechanisms dependent on dispersal, species interactions, trait evolution and geographic isolation processes. Customizable eco-evolutionary processes, which interact with dynamic landscapes, make it possible to adjust for various macro-eco-evolutionary hypotheses about specific taxonomic groups, ecosystem types or processes. We made the engine openly available to the research community in an R-package to catalyse an interdisciplinary exploration, application and quantification of the mechanisms behind biodiversity dynamics. The R statistical programming language and environment [65] is widely used for reproducible and open-source research, and since its origins it has been used for handling and analysing spatial data [66]. Gen3sis follows best practices for scientific computing [67], including high modularization; consistent naming, style and formatting; single and meaningful authoritative representation; automated workflows; version control; continuous integration; and extensive documentation.

Gen3sis operates over a grid-based landscape, either the entire globe or a specific region. The landscape used as input is defined by the shape of the colonizable habitat (e.g. land masses for terrestrial organisms), its environmental properties (e.g. temperature and aridity) and its connectivity to dispersal (e.g. the influence of barriers, such as rivers and oceans for terrestrial organisms). Gen3sis simulates species’ population range dynamics, traits, diversification and spatial biodiversity patterns in response to geological, biological and environmental drivers. Using a combined trait-based and biological species concept, gen3sis tracks the creation, dynamics and extinction of species ranges, which are composed by a set of sites occupied by species populations. Eco-evolutionary dynamics are driven by user-specified landscapes and processes, including ecology, dispersal, speciation and evolution (Figure 1). Below we explain the gen3sis inputs, the configurations (including eco-evolutionary processes), and the landscapes defining the computer model, as well as user-defined outputs (Figure 1 C–F)

**Figure 1.**
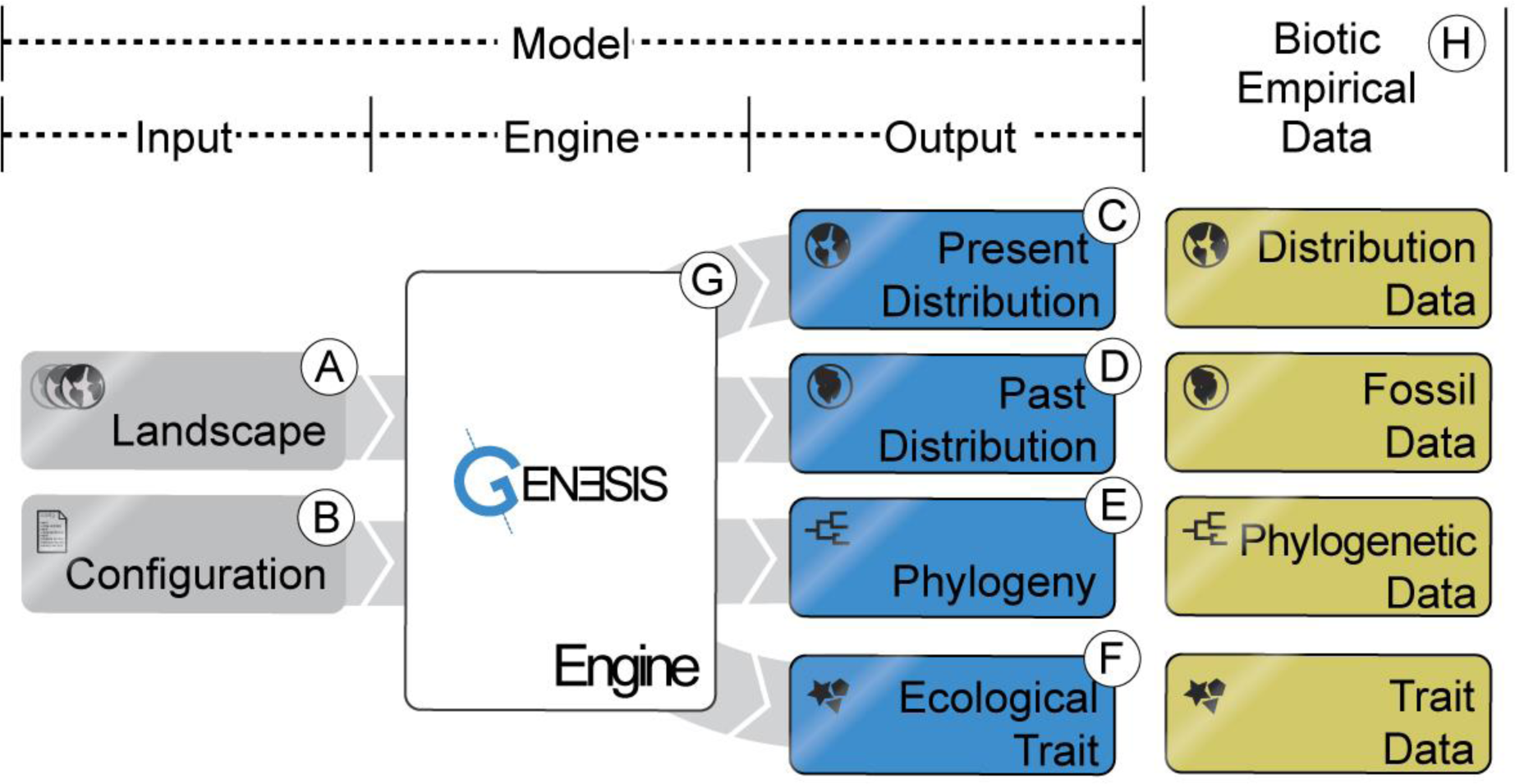
Schematic of the main components of the computer model: (A, B) model inputs, including the spatio-temporal landscape objects and the configuration file; (C–F) model outputs, including present and past species ranges, phylogenetic relationships among species, and the ecological traits of species; (G) model engine containing the mechanics; and (H) empirical data applicable for model validation.

### Inputs and initialization

Gen3sis has two input objects which define a particular model (Figure 1). These inputs are: (i) a dynamic landscape (Figure 1 A), which is further divided into environmental variables and distance matrices; and (ii) a configuration (Figure 1 B), in which the user can define initial conditions, biological functions and their parameter values, as well as technical settings for the model core.

#### Landscape

The landscape objects (Figure 1 A) form the spatio-temporal context in which the processes of speciation, dispersal, evolution and ecology take place. Landscape objects are generated based on temporal sequences of landscapes in the form of raster files, which are summarized in the form of two classes. The first landscape class contains: (i) the geographic coordinates of the landscape sites, (ii) the corresponding information on which sites are generally suitable for a clade (e.g. land or ocean), and (iii) the environmental conditions (e.g. temperature and aridity). The landscape may be simplified into a single geographic axis [e.g. 68] for theoretical experiments, or it may consider realistic configurations aimed at reproducing real local or global landscapes [24, 69, 70]. The second landscape class defines the connectivity of the landscape, used for computing dispersal and consequently isolation of populations. By default, the connection cost between occupied sites is computed for each time-step from the gridded landscape data based on haversine geographic distances. This can be modified by a user-defined cost function in order to account for barriers with different strengths (e.g. based on elevation [69], water or land) or even to facilitate dispersal in specific directions (e.g. to account for currents and river flow directions). The final connection costs are stored as sparse distance matrices [71]. Distance matrices, containing the connection costs, are provided at every time-step as either: (i) a pre-computed full distance matrix, containing all habitable sites in the landscape (faster simulations but more storage required); or (ii) a local distance matrix, computed from neighbouring site distances up to a user-defined range limit (slower simulations but less storage required).

#### Configuration

The configuration object (Figure 1 B) includes the customizable *initialization*, *observer*, *speciation*, *dispersal*, *evolution* and *ecology* functions. These six functions define a configuration applied in the simulation engine (Table 1). The possibility to customize these functions confers the high flexibility of gen3sis in terms of including a wide range of mechanisms, as illustrated by three configurations presented in a case study (Note S1, Table S1). Additionally, the configuration object lists the model settings, including: (i) whether a random seed is used, allowing simulation reproducibility; (ii) start and end times of the simulation; (iii) rules about aborting the simulation, including the maximum global or local species number allowed; and (iv) the list of ecological traits considered in the simulation. One or multiple traits can be defined, which should correspond to those used in the *ecology* function. Moreover, the *initialization* function creates the ancestor species at the start of the simulation. Users can define the number of ancestor species, their distribution within the ancient landscape and their initial trait values. With the *observer* function, changes over time in any abiotic or biotic information of the virtual world can be recorded by defining the outputs that are saved at specified time-steps. Outputs can be saved and plotted in real-time as the model runs. The core biological functions (i.e. *speciation*, *dispersal*, *evolution* and *ecology*) are presented below.

**Table 1.**
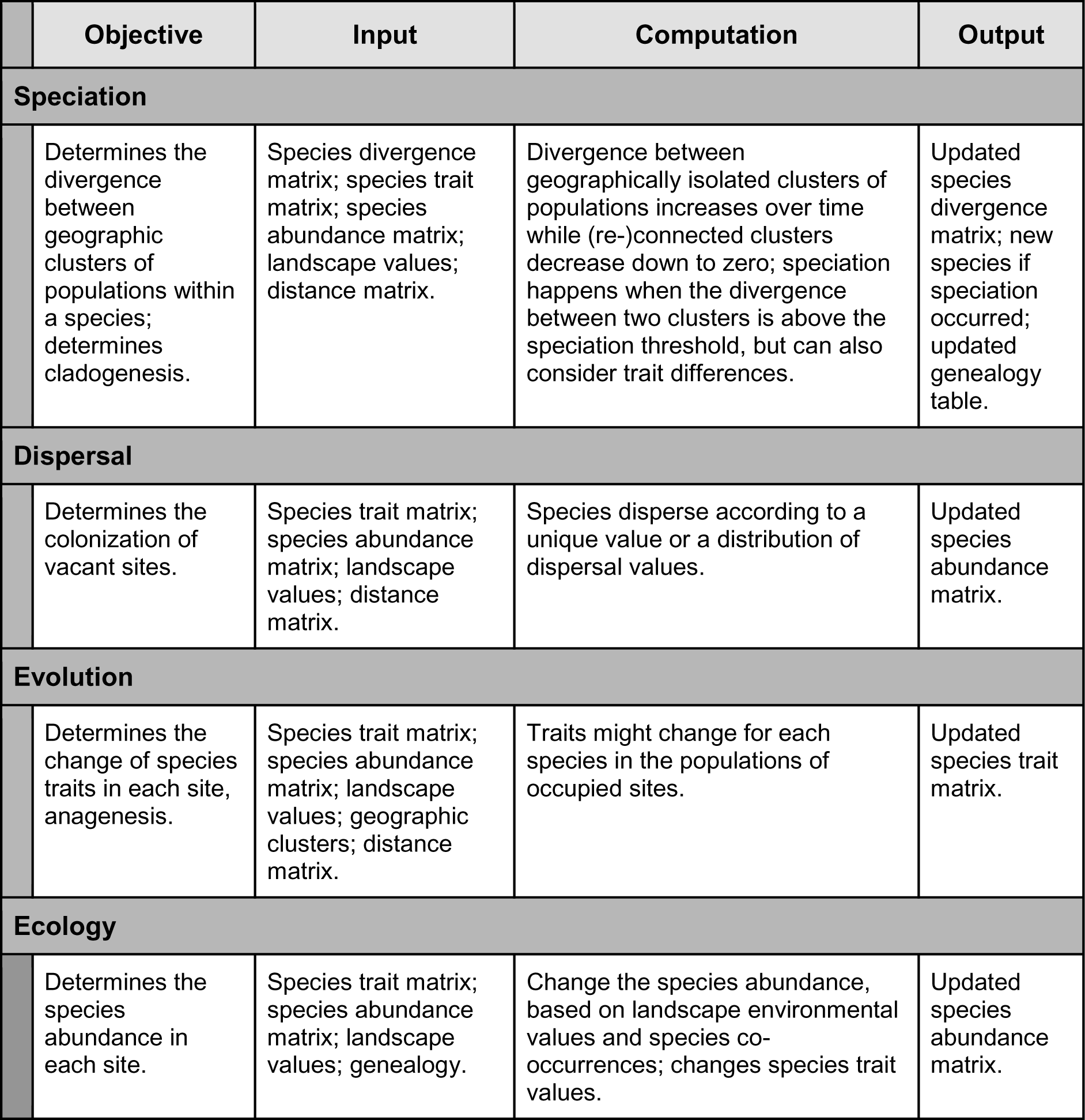
Presentation of the core functions of *speciation*, *dispersal*, *ecology* and *evolution* implemented in gen3sis. The computation of core functions is customizable in the configuration object. Shown are input objects that are combined to generate updated outputs. The table corresponds to the mechanisms presented in Figure 2 B.

#### Core functions and objects

The states of the computer model are updated in discrete time-steps. At each time-step, the *speciation*, *dispersal*, *evolution* and *ecology* functions are executed sequentially (Figure 2). Speciation and extinction emerge from interactions across core functions. For example, speciation events are influenced by *speciation* function as well as by the *ecology* and *dispersal* functions that interact in a dynamic landscape, ultimately dictating populations’ geographic isolation. Likewise, global extinctions depend on local extinctions, which are influenced by the *dispersal*, *evolution* and *ecology* functions that dictate adaptation and migration capacity. Internally, the computer model defines core objects of the simulations: species abundances; species trait values; the species divergence matrix between all populations for each species; and the phylogeny of all species created during the simulation. In the following sections, we describe the core processes in gen3sis, as well as their inputs and outputs. For a summary see Table 1.

**Figure 2.**
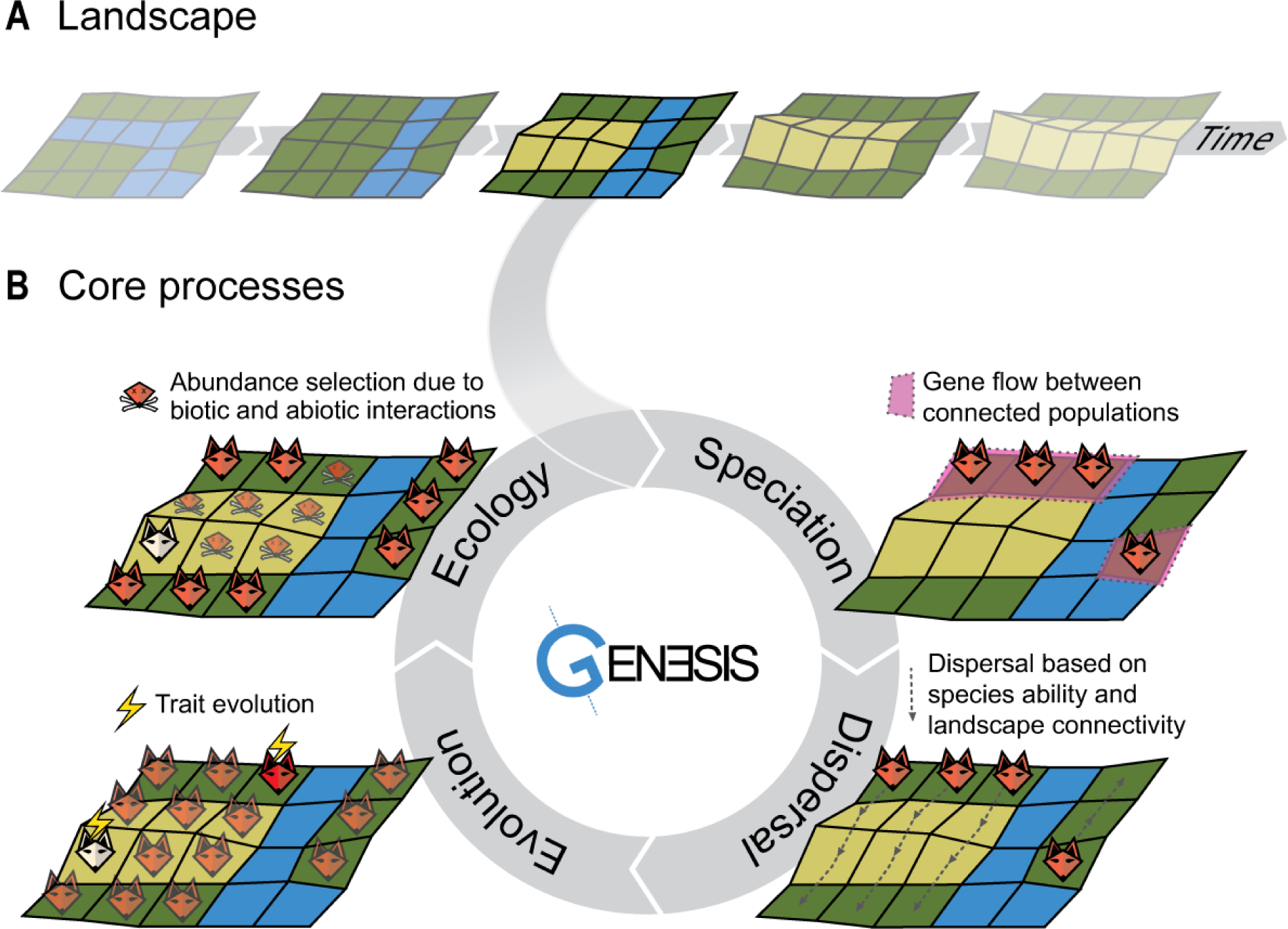
Schematic example of the gen3sis engine simulation cycle of one species’ populations over a landscape evolution example containing highlands (yellow), lowlands (green) and a river acting as a barrier (blue). (A) Landscape. A time series of landscapes is used as input, with the landscape being updated after every time-step of the simulation cycle, i.e. after the ecology process. (B) Model core processes. <u>First</u>, the speciation process determines the divergence between geographic clusters of populations that are not connected and splits the clusters into new species if a threshold is reached. In this illustration, divergence between clusters of fox populations was not sufficient to trigger speciation. <u>Second</u>, in the dispersal process, the species spreads within a landscape to reachable new sites. In this illustration, the river limits dispersal. <u>Third</u>, the evolution process can modify the value of the traits in the populations. In this illustration, two fox populations show trait evolution in their ability to cope with the local environment (i.e. red and white fox populations). <u>Fourth</u>, the ecology process recalculates the abundance of the species in each site based on the abiotic condition and co-occurring species, possibly resulting in local extinctions. In this illustration, the red fox was unsuited to the lowlands while the white fox survived in the highlands. Speciation and extinction events emerge from multiple simulation cycles of customizable processes.

Running a simulation in gen3sis consists of the following steps: (i) Read in the configuration object, prepare the output directories, load the initial landscape (Figure 2 A) and create the ancestor specie(s) (using the *initialization* function *create_ancestor_species*). (ii) Run the main loop over the landscape time-steps. At every time-step, the engine loads the appropriate landscape, removes all sites that became uninhabitable in the new time-step, and executes the core functions as defined by the configuration object (Figure 2 B). (iii) At the end of every time-step, gen3sis saves the species richness, genealogy and, if desired, the species, landscape and other customized observations that are defined in the *observer* function (e.g. summary statistics and species pattern plots). Core functions are modifiable and can account for a wide range of mechanisms, as illustrated in the case study (Notes S1 and S2). Conversely, functions can be turned off, for example in an ecologically neutral model. For a pseudo-code of gen3sis see Note S3.

### Speciation

#### Core

The *speciation* function iterates over every species separately, registers populations’ geographic occupancy (species range), and determines when geographic isolation between population clusters is sufficient to trigger a lineage-splitting event of cladogenesis. A species’ range can be segregated into spatially discontinuous geographic clusters of sites and is determined by multiple other processes. The clustering of occupied sites is based on the species’ dispersal capacity and the landscape connection costs. Over time, disconnected clusters gradually accumulate incompatibility (divergence), analogous to genetic differentiation. Disconnected species population clusters that maintain geographic isolation for a prolonged period of time will result in different species after the differentiation threshold Ϟ is reached (modelling Dobzhansky-Muller incompatibilities [72]). These clusters become two or more distinct species, and a divergence matrix reset follows. On the other hand, if geographic clusters come into secondary contact before the speciation occurs, they coalesce and incompatibilities are gradually reduced to zero.

#### Non-exhaustive modification possibilities

A customizable *speciation* function can further embrace processes that modulate speciation. Increased divergence values per time-step can be constant for all species or change depending on biotic and abiotic conditions, such as faster divergence between species occupying higher temperature sites [62], or they can be dependent on population size [73] or other attributes [74]. The function also takes the ecological traits as input, thus allowing for ecological speciation [22], where speciation depends on the divergence of ecological traits between – but not within – clusters [75].

### Dispersal

#### Core

The *dispersal* function iterates over all species populations and determines the connectivity between sites and the colonization of new sites in the grid cell. Dispersal distances are drawn following a user-defined dispersal function and then compared with the distance between pairs of occupied and unoccupied sites. A unique dispersal value can be used (deterministic connection of sites) or dispersal values can be selected from a specified distribution (stochastic connection of sites). If the occupied to unoccupied site connection cost is lower than the dispersal distance, the dispersal is successful. If populations in multiple origin sites manage to colonize an unoccupied site, a colonizer is selected randomly to seed the traits for the newly occupied site.

#### Non-exhaustive modification possibilities

A customizable *dispersal* function enables the modelling of different dispersal kernels depending on the type of organism considered. Dispersal values can be further linked with: the *ecology* function, for instance trade-off with other traits [76], e.g. dispersal versus competitive ability [77]; and the *evolution* function allowing dispersal to evolve, resulting in species with different dispersal abilities [78].

#### Evolution

##### Core

The *evolution* function determines the change in the traits of each population in occupied sites of each species. Traits are defined in the configuration object and can evolve over time for each species’ populations. The function iterates over every population of a species and modifies the trait(s) according to the specified function. Any number of traits, informed at the configuration object, can evolve (e.g. traits related to dispersal, niche or competition).

##### Non-exhaustive modification possibilities

A customizable *evolution* function takes as input the species abundance, species trait, species divergence clusters and the landscape values. In the function it is possible to define which traits evolve and how they change at each time-step. In particular, the frequency and/or amount of change can be made dependent on temperature [79], ecological traits [80], or abundances [81], while the directions of change can follow local optima or various evolutionary models, including Brownian motion [82] and Ornstein–Uhlenbeck [83].

#### Ecology

##### Core

The *ecology* function determines the abundance or presence of populations in occupied sites of each species. Thus, extinction processes derive from *ecology* function interactions with other processes and landscape dynamics. The function iterates over all occupied sites and updates the species population abundances or presences on the basis of local environmental values, updated co-occurrence patterns and species traits.

##### Non-exhaustive modification possibilities

A customizable *ecology* function takes as input the species abundance, species trait, species divergence and clusters, and the landscape values. Inspired by classic niche theory [10, 15, 84], the function can account for various niche mechanisms, from simple environmental limits to complex multi-species interactions. It is possible, for example, to include a carrying capacity for the total number of individuals or species [21] or competition between species based on phylogenetic or trait distances [26], based on an interaction currency [85], or further constrained by a functional trade-off [76].

### Outputs and comparisons with empirical data

The computer model delivers a wide range of outputs that can be compared with empirical data (Figure 1, Table 2). Gen3sis is therefore suitable for analysing the links between interacting processes and their multidimensional emergent patterns. By recording the time and origin of all speciation events, as well as trait distributions and abundance throughout evolutionary history, the simulation model records the information required to track the dynamics of diversity and the shaping of phylogenetic relationships. The most common patterns observed and studied by ecologists and evolutionary biologists, including species ranges, abundances and richness, are emergent properties of the modelled processes (Table 2). All internal objects are accessible to the observer function, which is configurable and executed during simulation runs. This provides direct simulation outputs in a format ready to be stored, analysed and compared with empirical data. Given the flexibility of gen3sis, it is possible to explore not only parameter ranges guided by prior knowledge available for a given taxonomic group, but also variations in landscape scenarios and mechanisms (Figure 3). Furthermore, validating modelled outputs with multiple empirical patterns is recommended [20, 23, 35]. Gen3sis generates multiple outputs, which can be compared with empirical data using simulation rankings or acceptance criteria [23, 35, 86].

**Figure 3.**
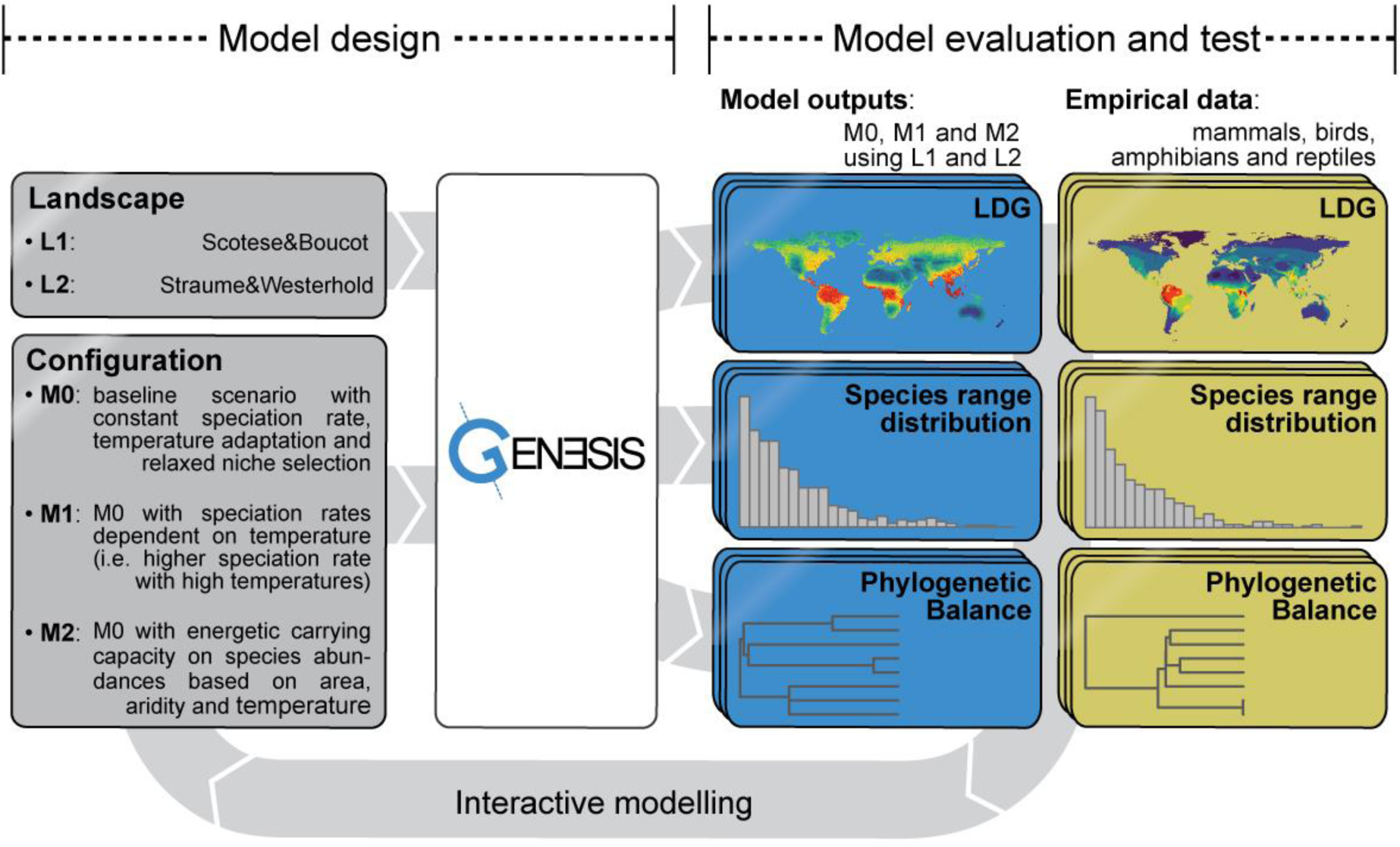
Schematic representation of the case study showing the model design with two landscapes (i.e. L1 and L2) and configurations of three models (i.e. M0, M1 and M2) (Table S1), and model evaluation and test, based on multiple patterns including: LDG, range size distributions and phylogenetic balance. Selection criteria were based on empirical data from major tetrapod groups, i.e. mammals, birds, amphibians and reptiles (Table 3).

**Table 2.**
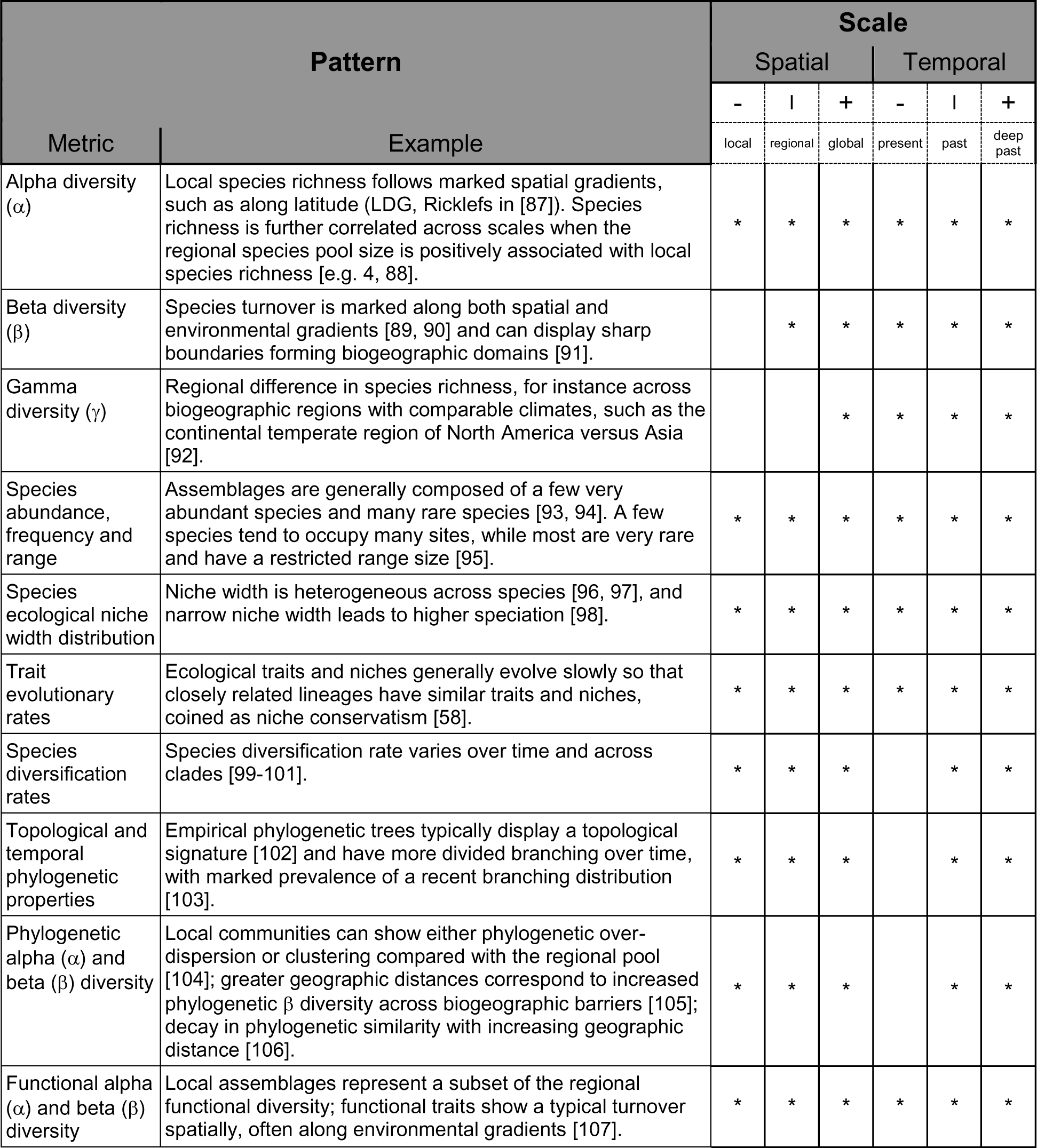
List of outputs from the gen3sis computer model, both direct and indirect, that can be compared with empirical data. Direct outputs are the species abundance matrix, species trait matrix and phylogeny, while indirect outputs result from various combinations of the direct outputs. The computations of indirect outputs rely on other packages available in the R environment [65].

**Table 3.** Model acceptance table with pattern descriptions and details of acceptance derived from empirical data. Percentages of accepted simulations (for both landscapes) are shown for each model and acceptance parameter and the combination of all acceptance patterns.

| Acceptance |  | M0 | M1 | M2 |
| --- | --- | --- | --- | --- |
| Pattern | Description and empirical acceptance | (n=480) | (n=720) | (n=480) |
| **LDG** | Percentage of species loss per latitudinal degree from linear regression slope.<br>Accept LDGs between 5% and 1% | 34% | 36% | 42% |
| **Phytogenic balance** | The imbalance of a phylogenetic tree is measured by the value that maximizes the likelihood in the β-splitting model [138].<br>Accept phylogenies with $\beta$ between -1.4 and -0.3 | 58% | 51% | 66% |
| **Range** | Range size distributions.<br>Accept only distributions that show a consistent frequency decrease towards large-ranged species with a tolerance of 5% | 0% | 0% | 5% |
| **Combined** | Simulations passing all criteria above with at least 100 species alive at the present. | 0% | 0% | 1% |

## Case study: The emergence of the LDG from environmental changes of the Cenozoic

### Context

The LDG is one of Earth’s most iconic biodiversity patterns, but the underlying mechanisms remain largely debated [20, 61, 62, 97, 98, 108–110]. Many hypotheses have been proposed to explain the formation of the LDG [20], and these generally agree that a combination of biological processes and landscape dynamics has shaped the emergence of the LDG [20]. Hypotheses can be generally grouped into three categories [20]: (i) *time for species accumulation*, (ii) variation in *diversification rates*, and (iii) variation in *ecological limits* [Table 1 in 20].

Tropical environments can be used to exemplify these three hypothesis categories: First, the times for species accumulation propose that since tropical environments are older, they should have more time for species accumulation, without assuming further specific ecological or evolutionary mechanisms [57–60]. Second, higher temperatures in the tropics increase metabolic and mutation rates, which could lead to faster reproductive incompatibilities among populations and higher speciation rates compared with colder environments [61, 62]. Third, the tropics are generally more productive than colder environments and greater resource availability can sustain higher abundances, and therefore a larger number of species can coexist there [63, 64, 111, 112].

We implemented one model for each of these hypotheses and simulated the spread, speciation, dispersal and extinction of terrestrial organisms over the Cenozoic. We evaluated whether the emerging patterns from these simulated mechanisms correspond to the empirical LDG, phylogenetic tree imbalance and range size frequencies computed from data of major tetrapod groups, including mammals, birds, amphibians and reptiles (Figure 3).

### Input landscapes

The Cenozoic (i.e. 65 Ma until the present) is considered key for the diversification of the current biota [113] and is the period during which the modern LDG is expected to have been formed [114]. In the Cenozoic, the continents assumed their modern geographic configuration [24]. Climatically, this period was characterized by a general cooling, especially in the Miocene, and ended with the climatic oscillations of the Quaternary [115]. We compiled two global paleoenvironmental landscapes (i.e. L1 and L2) for the Cenozoic at 1° and ∼170 kyr of spatial and temporal resolution, respectively (Note S1, Animations S1 and S2). To account for uncertainties on paleo-reconstructions on the emerging large-scale biodiversity patterns, we used two paleo-elevation reconstructions [116, 117] associated with two approaches to estimate the paleo-temperature of sites (Note S1). L1 had temperatures defined by Köppen bands based on the geographic distribution of lithologic indicators of climate [54]. L2 had temperature defined by a composite of benthic foraminifer isotope records over time [118] and along latitude for specific time periods [119–125]. An aridity index ranging from zero to one was computed based on the subtropical arid Köppen zone for both landscapes [54]. For details see Note S1.

### Hypothesis implementation

We implemented three hypotheses explaining the emergence of the LDG as different gen3sis models. The models (i.e. M0, M1 and M2) had distinct speciation and ecological processes (Figure 3, Note S1, Table S1). All simulations were initiated with one single ancestor species spread over the entire terrestrial surface of the Earth at 65 Ma, where the temperature optimum of each population matched local site conditions. Since we focused on terrestrial organisms, aquatic sites were considered inhabitable and twice as difficult to cross as terrestrial sites. This approximates the different dispersal limitation imposed by aquatic and terrestrial sites. The spherical shape of the Earth was accounted for in distance calculations by using haversine geodesic distances. Species disperse following a Weibull distribution with shape 2 or 5 and a scale of 550, 650, 750 or 850, resulting in most values being around 500–1500 km, with rare large dispersal events above 2000 km. The *evolution* function defines the temperature niche optimum to evolve following Brownian motion. Temperature niche optima are homogenized per geographic cluster by an abundance-weighted mean after ecological processes happen. We explored three rates of niche evolution, with a standard deviation equivalent to ±0.1°C, ±0.5°C and ±1°C.

#### M0

In the implementation of the *time for species accumulation*, the *ecology* function defines the species population abundance, where the abundance increases proportionally to the distance between the population temperature niche optimum and the site temperature (Note S1). Clusters of populations that accumulated differentiation over Ϟ = 12, 24, 36, 48 and 60 will speciate, corresponding to events occurring after 2, 4, 6, 8 and 10 myr of isolation, respectively. The divergence rate between isolated clusters was kept constant (i.e. +1 for every 170 kyr of isolation). Model M0, assuming *time for species accumulation*, acted as a baseline model. This means that all mechanisms present in this model were the same for M1 and M2 if not specified otherwise.

#### M1

In the implementation of the *diversification rates,* the speciation function applies a temperature-dependent divergence between population clusters [61, 62]. Species in warmer environments accumulate divergence between disconnected clusters of populations at a higher rate (Note S1). The rate of differentiation increase was the average site temperature of the species clusters to the power of 2, 4 or 6 plus a constant. This created a differentiation increase of +1.5 for isolated clusters of a species at the warmest range and +0.5 at the coldest range for every 170 kyr of isolation (Note S1, Figure S1). Using Ϟ = 12, 24, 36, 48 and 60, this corresponds to a speciation event after 1.3, 2.7, 4.0, 5.3, 6.7 myr and after 4, 8, 12, 16, 20 myr for the warmest and coldest species, respectively.

#### M2

In the implementation of the *ecological limits*, the *ecology* function includes a carrying capacity *k* of each site that scales with area energy and aridity in the ecology function [112, 126]. The theory of carrying capacity proposes that energy limits abundances and therefore determines how many of each species can coexist in a given place [21, 112]. If the sum of all species abundances in a site is above *k*, species abundances are randomly reduced across species until *k* is reached. We explored low and high *k* values using a *k* power-law scaling of 2 and 3.

### Exploration of model parameters

For each model (i.e. M0, M1 and M2) in combination with each landscape (i.e. L1 and L2), we explored a range of conservative model parameters in an interactive modelling cycle (Figure 3). In addition, we explored dispersal distributions and parameters ranging in realized mean and 95% quantiles between less than a single cell, i.e. ∼50 km for a landscape at 4°, and more than the Earth’s diameter, i.e. ∼12’742 km (Figure S2). Trait evolution frequency and intensity ranged from zero to one. We ran a full factorial exploration of these parameter ranges at a coarse resolution of 4° (i.e. M0 n=480, M1 n=720, M2 n=480) and compared these to empirical data. Simulations considered further: (i) had at least one speciation event; (ii) did not have all species becoming extinct; (iii) had fewer than 50’000 species; or (iv) had fewer than 10’000 species cohabiting the same site at any point in time (Note S1). After parameter range exploration, we identified realistic parameters and ran a subset at 1° for high-resolution outputs (Figure 4).

**Figure 4.**
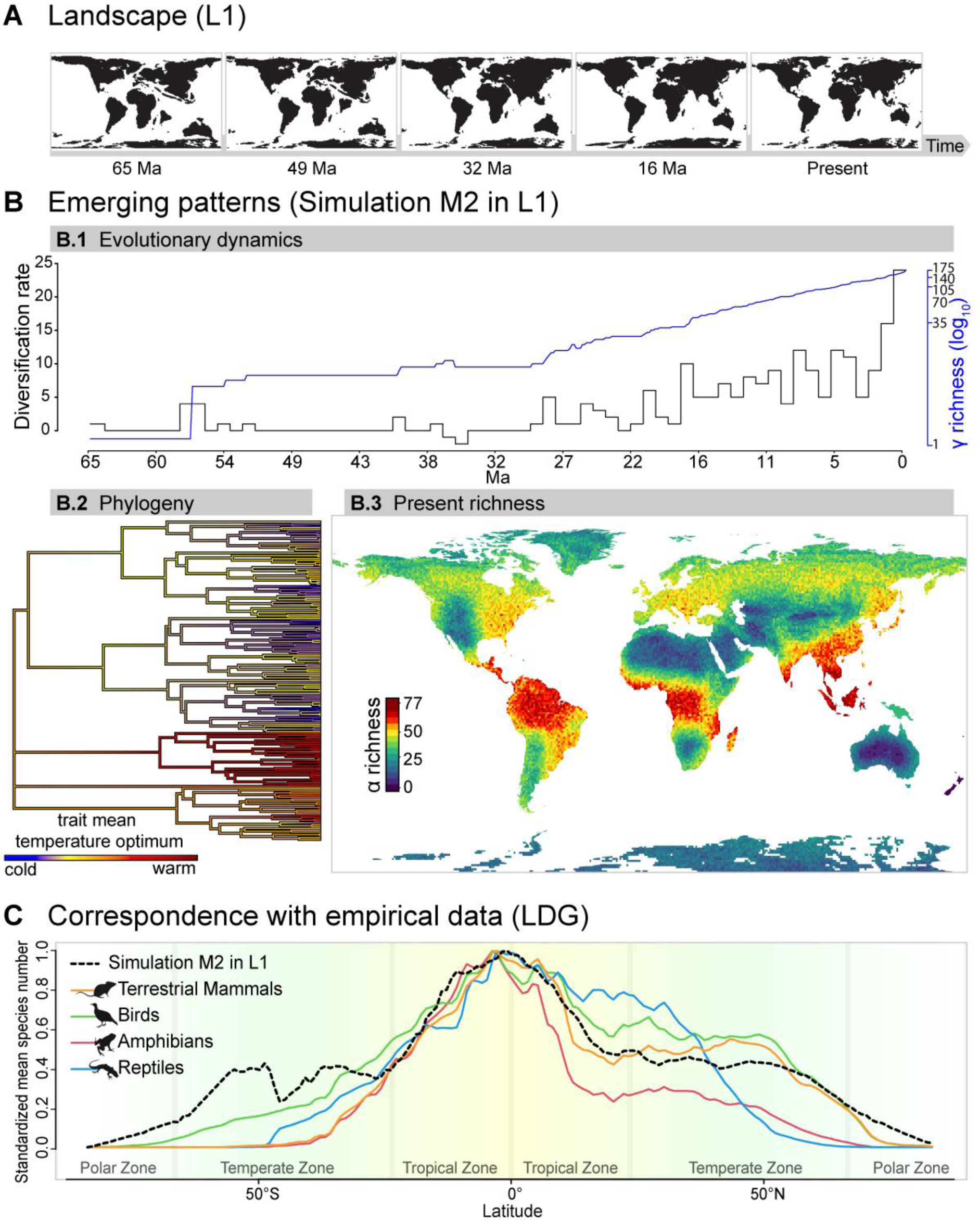
Illustration of one global simulation of the speciation, dispersal and extinction of lineages over the Cenozoic, starting with a single ancestor species and imposed energetic carrying capacity (M2 in L1). (A) Images of the Earth land masses through time, used as input for the simulation. (B) Selected emerging patterns: evolutionary dynamics; phylogeny; and present richness. (B.1) Evolutionary dynamics: γ richness (log10 scale) through time (blue line) and diversification rate. (B.2) Phylogeny showing the distribution of the temperature optima for all extant species. (B.3) Present distribution of simulated α biodiversity globally, which indicates locations of biodiversity hotspots. For the empirical match see Figure S3. (C) Model correspondence with empirical data of terrestrial mammals, birds, amphibians and reptiles for the LDG, measured as the standardized and area-scaled mean species number per latitudinal degree.

### Correspondence with empirical data

In order to explore the parameters of all three models and compare their ability to produce the observed biodiversity patterns, we used a pattern-oriented modelling (POM) approach [23, 86]. POM compares the predictions of each model and parameter combination with a number of diagnostic patterns from empirical observations. In our case, we used the LDG slope, tree imbalance and range size frequencies as diagnostics patterns (Figure 3, Note S1). The POM approach allows a calibration and model comparison based on high-level diagnostic patterns, avoiding the hurdles of defining explicit (approximate) likelihood functions [127]. The POM approach requires the specification of a range for each pattern under which observation and prediction are accepted, hence when a simulation satisfactorily reproduces empirical observations. Unless POM is coupled with an explicit probabilistic model [127], the limits for acceptance must be decided by the modeller based on their understanding of the data [23, 86].

To generate the empirical values for these patterns, we obtained distribution data on 25’941 species [128–130], following [131], and phylogenetic data on 18’978 species [5, 132–135], following [136] for major tetrapod groups, i.e. terrestrial mammals, birds, amphibians and reptiles (Note S1). LDG is given by the percentage of species loss per latitudinal degree and measured by the slope of a linear regression on normalized species richness against absolute latitude. β-statistics [31] was used for phylogenetic tree imbalance in ultrametic trees, following [102]. Species ranges decrease (SRD) in km^2^ is given by the percentage of species loss per species range and is measured by the slope of a linear regression of range size distributions. Empirical values of LDG, β and SRD were: mammals (LDG=5.1%, β=-0.4, SRD=2.3*10^3^%), birds (LDG=1.5%, β=-1.3, SRD=6.5*10^7^%), amphibians (LDG=3.9%, β=-0.7, SRD=0.11%) and reptiles (LDG=1.5%, β=-0.8, SRD=5.3*10^3^%). Based on these values, we used the following acceptance criteria: (i) LDG between 5.4% and 1.1%, (ii) tree shape statistic β between -1.4 and -0.3, and (iii) range size frequencies with a decrease in the number of large-range species with a tolerance of 5% [93–95] (Note S1).

### Simulations results and synthesis

We found that model M2 was the best match for all the empirical patterns individually, and the only model able to pass all acceptance criteria (Table 3). Although all three models were able to reproduce the LDG, M2 was superior in explaining the LDG, phylogenetic tree imbalance and species range size frequencies simultaneously (Table 3). Most simulations of model M2 (67%) resulted in a decrease in species richness at higher latitudes, indicating that the LDG emerged systematically under M2 mechanisms (Figure S3, Tables S2, S3 and S4). Increasing the spatial resolution of the simulations (n=12) resulted in an increase in γ richness and computation time and a slight decrease of the LDG (Figure S5), which was associated with adisproportionally larger number of sites towards higher latitudes, which also affects population connectivity and therefore speciation rates [137]. We then selected the best matching simulation of M2 in L1 at 1° (n=12) that predicted realistic biodiversity patterns (Figure 4, Animation S4), The emerging LDG (i.e. 4.6% of species loss per latitudinal degree) closely matched empirical curves, with good agreement for mammals (Pearson r=0.6), birds (r=0.57), amphibians (r=0.57) and reptiles (r=0.38) (Note S1, Figure 4C, Figure S6). Finally, we found that the support for M2 over M0 and M1 was consistent across the two alternative landscapes L1 and L2 (Figure S3, Table S4).

**Table 4.** A non-exhaustive list of expected applications of gen3sis. Given the flexibility and the range of outputs produced by the engine, we expect that gen3sis will serve a large range of purposes, from testing a variety of theories and hypotheses to evaluating phylogenetic diversification methods

| Use | Examples from Figure 1 |
| --- | --- |
| Testing phylogenetic inference methods, including diversification rates in phylogeographic reconstructions. | Infer diversification rate in gen3sis simulated phylogenies (E) and compare with a known diversification in gen3sis (A, B & G). |
| Providing biotic scenarios for past responses to geodynamics. | Based on model outputs (C–F) and comparisons with empirical data (H), select plausible models (B). |
| Testing paleo-climatic and paleo-topographic reconstructions using biodiversity data. | Based on model outputs (C–F) and comparisons with empirical data (H), select plausible landscape(s) (A). |
| Comparing expectations of different processes relating to the origin of biodiversity; generating and testing hypotheses. | Compare models (A, B & G) with outputs (C–F) and possibly how well outputs match empirical data (H). |
| Comparing simulated intra-specific population structure with empirical genetic data. | Compare simulated divergence matrices with population genetic data. |
| Forecasting the response of biodiversity to global changes (e.g. climate or fragmentation). | Extrapolate plausible and validated models (A, B & G) on landscapes under climate change scenarios (A). |
| Investigating trait evolution through space and time. | Combine past and present simulated species traits (F) and distributions (C, D) with fossil and trait data (H). |
| Modelling complex systems in space and time in unconventional biological contexts in order to investigate eco-evolutionary processes in fields traditionally not relying on biological principles. | Model eco-evolutionary mechanisms (A, B & G) in an unconventional eco-evolutionary context. |

Our sensitivity analyses of parameters further provided information about the role of dispersal and ecological processes in shaping the LDG (Note S1, Table S2 and S3). In particular, our results indicate that an increase in the scaling factor of carrying capacity with energy k resulted in a steeper LDG slope, which is in agreement with findings from previous studies [21, 61, 112, 126]. Similarly, increasing the time for divergence consistently led to lower species richness and flattened the LDG slope so that the tropics accumulated diversity more slowly, but changes in speciation rates were less likely to drive large-scale biodiversity patterns [110]. Saupe and colleagues [25] showed that simulations with poor dispersal are better at representing the observed strong LDG in tetrapods. In agreement with their results, our parameter explorations indicated that dispersal correlated negatively with LDG [25], and simulations with lower dispersal parameters agreed better with the data (Note S1). While previous case studies [25, 26, 44] have been carried out to investigate the formation of the LDG using computer models, they used a shorter timeframe (i.e. below 1 Ma) and/or explored few mechanisms, i.e. simplified landscape or single acceptance criteria [24, 41, 42, 110]. Beyond this illustrative case study, future analyses could combine multiple mechanisms in relation to additional biodiversity patterns in order to investigate the most likely combination of mechanisms shaping the intriguing LDG pattern.

## Discussion

Understanding the emergence of biodiversity patterns requires the consideration of multiple biological processes and abiotic forces that potentially underpin them [20, 26, 35, 36]. We have introduced gen3sis, a modular, spatially-explicit, eco-evolutionary simulation engine implemented as an R-package, which offers the possibility to explore ecological and macroevolutionary dynamics over changing landscapes. Gen3sis generates commonly observed diversity patterns and, thanks to its flexibility, enables the testing of a broad range of hypotheses (Table 4). It follows the principle of computer models from other fields [139–141], where mechanisms are implemented in a controlled numeric environment and emerging patterns can be compared with empirical data [23]. The combination of exploring patterns emerging from models and matching qualitatively and quantitatively the model outputs to empirical data should increase our understanding of the processes underlying global biodiversity patterns.

Using a case study, we have illustrated the flexibility and utility of gen3sis in modelling multiple eco-evolutionary hypotheses in global paleo-environmental reconstructions (Figures 3 and 4). Our findings suggest that global biodiversity patterns can be modelled realistically by combining paleo-environmental reconstructions with eco-evolutionary processes, thus moving beyond pattern description to pattern reproduction [35]. Nevertheless, in our case study we only implemented a few of the standing LDG hypotheses [20, 34]. Multiple macroecological and macroevolutionary hypotheses still have to be tested, including the role of stronger biotic interactions in the tropics than in other regions [142], and compared with more biodiversity patterns [20]. Considering multiple additional biodiversity patterns will allow a more robust selection of models. Apart from the global LDG case study, we propose an additional case study (Note S2, Figure S7) illustrating how gen3sis can be used for regional and theoretical studies, such as investigations of the effect of island ontology on the temporal dynamics of biodiversity [39, 143]. Further, illustrations associated with the programming code are offered as a vignette of the R-package, which will support broad application of gen3sis. Altogether, our examples illustrate the great potential for exploration provided by gen3sis, promising future advances in our understanding of empirical biodiversity patterns.

Verbal explanations of the main principles underlying the emergence of biodiversity are frequently proposed but are rarely quantified or readily generalized across study systems [20]. We anticipate that gen3sis will be particularly useful for exploring the consequences of mechanisms that so far have mostly been verbally defined. For example, the origins of biodiversity gradients have been associated with a variety of mechanisms [7], but these represent verbal abstractions of biological processes that are difficult to evaluate [20]. Whereas simulation models can always be improved, their formulation implies formalizing process-based abstractions via mechanisms expected to shape the emergent properties of a system [144]. Specifically, when conveying models with gen3sis, decisions regarding the biological processes and landscapes must be formalized in a reproducible fashion. By introducing gen3sis, we encourage a standardization of configuration and landscape objects, which will facilitate future model comparisons. This standardization offers a robust framework for developing, testing, comparing, and applying the mechanisms relevant to biodiversity research.

Studying multiple patterns is a promising approach in disentangling competing hypotheses [20, 86]. A wide range of biodiversity dimensions can be simulated with gen3sis (Table 2), which – after appropriate sampling [145] – can serve in a multi-dimensional comparison with empirical data, i.e. a time series of species abundance matrices and trait matrices, as well as a phylogeny. These output objects are compatible with most R-packages used for community or phylogenetic analyses. Hence, the model outputs can be linked to packages computing diversification rates [146], community phylogenetics [147], or functional diversity [148]. The comparison of simulation outputs with empirical data requires a systematic exploration of processes, when formulating models, and parameter values [e.g. 149]. First, a set of mechanisms and/or a range of reasonable parameter values are explored, for instance dispersal distances from measurements in a specific clade [150] and/or evolutionary rates [151]. A range of simulation outputs can then be evaluated quantitatively by studying the range of models and parameter values that produce the highest level of agreement with multiple types of empirical data, using for example a pattern-oriented modelling approach [86]. For each model, patterns are evaluated given an acceptance criteria [e.g. 40]. A multi-scale and multi-pattern comparison of simulations with empirical data can be completed to evaluate a model’s ability to simultaneously reproduce not only one, but a diverse set of empirical patterns across multiple biodiversity dimensions.

The quality of the outputs of simulation models such as gen3sis hinges on accurate reconstructions of past environmental conditions [3, 42]. Although recent studies using realistic landscapes and computer models reproduced biodiversity patterns over a time scale spanning the Quaternary [25, 26, 44], many speciation and extinction events shaping present diversity patterns date back before the glaciation, and few studies have covered deep-time dynamics [24, 41, 42, 131]. Deep-time landscape reconstructions are still generally lacking but are increasingly becoming available [116, 118]. Here, we used available paleo-elevation models [116, 117] and paleoclimate indicators [54, 118–125, 152–154] to generate input landscapes to explore the formation of the LDG and account for uncertainties and limitations. For instance, we represented Quaternary climatic oscillation using ∼170 kyr time-steps, which correspond to a coarser temporal scale compared with the frequency of oscillations, and thus do not account for shorter climatic variation effects on diversity patterns [25, 26, 44]. We also did not consider ice cover, that can mask species’ habitable sites, which probably explains the the mismatch between simulated and empirical LDG patterns below 50° (Figure 4C). Moreover, paleo indicators of climate from Köppen bands have major limitations, and the temperature estimation derived in our case study can suffer from large inaccuracies. Lastly, extrapolation of the current temperature lapse rate along elevation might lead to erroneous estimates, especially in terms of the interaction with air moisture [155], which was not further investigated here. Hence, the presented case study represents a preliminary attempt for illustrative purposes. Further research is required to generate more accurate paleolandscapes, and research in biology should improve empirical evidence and our understanding of mechanisms. We expect that gen3sis will support exciting interdisciplinary research across the fields of geology, climatology and biology to understand the shaping of biodiversity.

## Conclusions

Here we have introduced gen3sis, a modular simulation engine that enables exploration of the consequences of ecological and evolutionary processes and feedbacks on the emergence of spatio-temporal macro-eco-evolutionary biodiversity dynamics. This modelling approach bears similarity with other computer models that have led to significant progress in other fields, such as climatology [139], cosmology [140] and conservation [141]. We showcase the versatility and utility of gen3sis by comparing the ability of three alternative mechanisms in two landscapes to generate the LDG while accounting for other global biodiversity patterns. Besides the LDG, frontiers on the origins of biodiversity involve [16]: (i) quantifying speciation, extinction and dispersal events [114]; (ii) exploring adaptive niche evolution [26, 44]; and (iii) investigating multiple diversity-dependence and carrying capacity mechanisms [21, 111, 112]. Further exploration possibilities may include: (iv) revealing the mechanisms behind age-dependent speciation and extinction patterns [102, 108, 156]; (v) contrasts between terrestrial and aquatic ecosystems [16]; and (vi) calculations of uncertainty resulting from climatic and geological dynamics [e.g. 24, 25, 26, 41, 42]. Gen3sis can support these research frontiers as a general tool for formalizing and studying existing theories associated with the origin of biodiversity, testing new hypotheses against data, and making predictions about future biodiversity trajectories (Table 4). Openly available as an R-package, gen3sis has the potential to catalyse interdisciplinary biodiversity research. We call for the formation of a community of ecologists, biologists, mathematicians, geologists, climatologists and scientists from other fields around this class of eco-evolutionary simulation models in order to unravel the processes that have shaped Earth’s biodiversity.

## Availability

Gen3sis is implemented in a mix of R and C++ code, and wrapped into an R-package. All high-level functions that the user may interact with are written in R, and are documented via the standard R / Roxygen help files for R-packages. Runtime-critical functions are implemented in C++ and coupled to R via the Rcpp framework. Additionally, the package provides several convenience functions to generate input data, configuration files and plots, as well as tutorials in the form of vignettes that illustrate how to declare models and run simulations. The software, under an open and free GPL3 license, can be downloaded from CRAN at https://CRAN.R-project.org/package=gen3sis. The development version, open to issue reporting and feature suggestions, is available at https://github.com/project-Gen3sis/R-package. Supporting information, such as notes, scritps, data, figures and animations, are available at https://github.com/ohagen/SupplementaryInformationGen3sis, facilitating full reproducibility.

## Acknowledgements

We thank Samuel Bickel and Alex Skeels for thorough comments on this manuscript and package. We thank Camille Albouy, Charles N.D. Santana, Lydian Boschman, Wilhelmine Bach, Thomas Keggin, Flurin Leugger, Victor L.J. Boussange, Conor Waldock and all sELDiG working group participants for insightful feedback during the model development. We thank the WSL and ETH Zürich for support and infrastructure including access to High Performance Computing facilities.

## Supporting Information captions

### Animations

**Animation S1** Reconstructed dynamic landscape L1 (i.e. world 65 Ma) with the environmental values used for the main case study.

**Animation S2** Reconstructed dynamic landscape L2 (i.e. world 65 Ma) with the environmental values used for the main case study.

**Animation S3** Theoretical dynamic landscape (i.e. theoretical island) with the environmental values used for the supplementary case study.

**Animation S4** Dynamic simulated biodiversity patterns (i.e. M2 L1 world from 65 Ma to the present). The map shows the α diversity and the top and right graphs indicate the richness profile of longitude and latitude, respectively.

### Figures

**Figure S1** Divergence increase per time-step *d_i_* against the normalized occupied niche of isolated populations for models (A) M0 and M2, which assume temperature-independent divergence; and (B) M1, which assumes temperature-dependent divergence, where divergence relates to the mean of the realized temperature with three different *d_power_* values.

**Figure S2** Non-exhaustive probability density functions of the explored dispersal parameters in a Weibull distribution with shape ɸ of 1, 2 and 5 and Ψ of 550, 650, 750 and 850.

**Figure S3** Frequencies of simulated normalized LDG slope (histogram) with empirical LDG for four main groups (dashed grey line) and acceptance range (red line). Frequencies for *models (A) M0, (B) M1, (C) M2 with total frequency and frequency discriminated for each landscape, i.e. L1 and L2*.

**Figure S4** Correlation of model parameters and three emerging patterns for all models and landscapes (A) M0 L1, (B) M0 L2, (C) M1 L1, (D) M1 L2, (E) M2 L1, and (F) M2 L2. *Emerging patterns: (i) <u>phylogeny beta</u> is the phylogenetic tree imbalance statistic measured as the value that maximizes the likelihood in the β-splitting model; (ii) <u>range quant 0.95%</u> is the value of the 95% quantile of the species range area distribution and; (iii) <u>LDG slope</u> is the slope of the linear regression of species richness*.

**Figure S5** Effects of grid cell size on simulations of M2 L1. (A) Correlation of grid cell, LDG slope and other summary statistics. (B) Simulated LDG slope and grid cell size, showing a significant effect of spatial resolution on LDG slope.

**Figure S6** Normalized richness of (A) selected simulation, (B) terrestrial mammals, (C) birds, *(D) amphibians and (E) reptiles, with Pearson correlation values for comparisons between simulated and empirical data*.

**Figure S7** Results of the island case study showing (A) landscape size and environmental dynamics and (B) results of three experiments (i.e. lower, equal and higher trait evolution compared with the temporal environmental variation). The time series in (B) shows γ richness (log10 scale) on theoretical oceanic islands, following the geomorphological dynamics of islands. Thick lines indicate the average of the replicates, whereas thin lines indicate SD envelopes (n=30 for each trait evolutionary rate scenario). The dashed grey vertical bar crossing the entire plot indicates the period in which the island reaches its maximum size.

### Notes

**Note S1** Global case study: emergence of the LDG from environmental changes of the Cenozoic.

**Note S2** Island case study: does trait evolution impact biodiversity dynamics?

**Note S3** Gen3sis pseudo-code.

